# Evolutionary history of the Galápagos Rail revealed by ancient mitogenomes and modern samples

**DOI:** 10.1101/2020.10.07.326983

**Authors:** Jaime A. Chaves, Pedro J. Martinez-Torres, Emiliano A. Depino, Sebastian Espinoza-Ulloa, Jefferson García-Loor, Annabel Beichman, Martin Stervander

## Abstract

The biotas of the Galápagos Islands are probably one of the best studied island systems and have provided a broad model of insular species’ origins and evolution. Nevertheless, some Galápagos species remain poorly characterized, such as the Galápagos Rail *Laterallus spilonota*. This bird species is one of the less explored groups of endemic vertebrates on these islands, due to its elusive behavior, cryptic plumage and restricted distribution. To date there is no genetic assessment of its origins and sister relationships to other taxa, and importantly, there is no data on its current genetic diversity. This lack of information is critical given the adverse fate of island rail species around the world in the recent past. Here we examine the genetics of Galápagos Rails using a combination of mitogenome *de novo* assembly with multi-locus sequencing (mtDNA+nuDNA) from both modern and historical samples. We show that the Galápagos Rail is part of the ‘American black rail clade’, sister to Black Rail *L. jamaicensis*, with a colonization of Galápagos dated to 1.2 Mya. The separate analysis of cyt*b*, ND2, and RAG-1 markers demonstrates shallow population structure across sampled islands, possibly due to elevated island connectivity. Additionally, birds sampled from Pinta possessed the lowest levels of genetic diversity, most likely reflecting the impact of past bottlenecks due to habitat loss caused by invasive goats grazing on sensitive habitat. The data presented here highlights the low genetic diversity in this endemic rail species and suggests the use of genetic data (both modern and historical) to guide conservation efforts.

## 1. Introduction

Studies of island biotas provide insights into the origin of species and the associated factors promoting it. From characterizing morphological variation, documenting the time of divergence from mainland counterparts, to ultimately understanding the formation of independently evolving lineages, insular species have inspired scientists for centuries [1–6]. One such biological assemblage are species found on the Galápagos archipelago. Generalities about Galápagos fauna have for example supported the hypothesis of island colonization in relation to island age (i.e. progression rule) [7,8], provided text book examples of the origin of adaptive radiations (i.e. Darwin’s finches) [3,9], postulated the role of isolation in promoting diversification (i.e. mockingbirds *Mimus* spp. [10], giant tortoises *Chelonoidis* spp. [11]), and reported various origins for endemic species [12,13]. Most of these studies focus on highly charismatic species, leaving almost no conspicuous species on the Galápagos unstudied. In particular, endemic vertebrate species have been subject to extensive molecular and morphological assessment providing one of the greatest insular biota datasets. The data has helped to raise awareness of conservation status and proposed management recommendations for the preservation of several of these species. Unfortunately, though, a few less charismatic endemic vertebrate species have suffered from a lack of such effort. This has not only left a gap in our understanding of their evolutionary trajectory and origin, but most importantly, their conservation status is poorly known and their extinction could potentially go unnoticed.

Species inhabiting oceanic islands are the most vulnerable to extinction due to human impact, particularly those that are flightless. Flightlessness in birds has evolved as a consequence of the absence of natural predators [14–16] creating opportunity for endemism. Rails (Rallidae) present high levels of endemism, in many cases restricted to a single island [17,18]. Unfortunately, the introduction of predators to such isolated regions make these species extremely susceptible to population declines and ultimately extinction [19,20]. Historically, rails have succumbed quickly to human contact with the loss of as many as 440–1,580 species on islands in the Pacific [21], where 22 of the 33 currently threatened rail species (World Conservation Union) occur on islands, of which 86% are threatened by invasive mammals (BirdLife International - IUCN databases).

The endemic Galápagos Rail *Laterallus spilonota* is an example of both this extreme adaptation to an isolated system of oceanic islands and the fate of inconspicuousness, being the least studied land-bird species on the Galápagos Islands. The historical distribution of Galápagos Rails has been documented by collectors and naturalists since the early 1900s, allowing a reconstruction of the impact and decline of populations compared to present data [22]. The introduction of rats and goats in the 18^th^ century, by mariners and early colonists using these islands for water and food supply [23–25], has had a direct impact on native species’ survival and ecosystem modification [26]. Galápagos Rails, once abundant as reported by Darwin (1896), depend on the presence of wetlands and dense vegetation, but these habitats were decimated and eroded by grazing goats and agricultural expansion, and altered by invasive plant species [27,28]. Likewise, rats and cats have had a negative impact on the survival of rails given their inability to fly [29]. These events have resulted in the extinction of several of the Galápagos Rail populations across the archipelago and a dramatic decline in surviving populations [30,31]. Galápagos Rails are currently restricted to small pockets of natural habitat in the highlands on five of the eight islands they historically inhabited. The species is currently listed as Vulnerable (Red Data List)[32] and despite the eradication of goats since the 1970s and the ongoing pest control efforts [33–35], rails still face continuous threats of habitat modification by invasive species [36] and expansion of agriculture [37].

Galápagos Rails are one of the least studied land bird species on the Galápagos: i) there is no genetic assessment of its phylogenetic relationships to other rails, ii) there are no estimations of the time since its colonization, iii) no inference of its phylogeographic patterns and inter-island genetic relationships, and importantly iv) there is no assessment of its current genetic diversity. This last effort is critical if we acknowledge the adverse fate of rails around the world in the recent past and thus, the uncertainty of its evolutionary potential in the future.

Here, we focus on alleviating these aspects and bringing to light the evolutionary history of this enigmatic endemic land bird of the Galápagos. We present data from a combination of fresh tissue samples and century-old historic museum specimens collected by the California Academy of Sciences expedition to the Galápagos in 1905–1906. The phylogenetic relationships for the Galápagos Rail proposed here for the first time are based on high-throughput sequenced *de novo* assembly of its mitochondrial genome (mitogenome), placing this species into a phylogenetic context. We also infer the timing of long-distance colonization and focus on genetic diversity and relationships between islands.

## 2. Materials and Methods

### 2.1. Study sites, sampling, and morphological data

We focused the large-scale phylogenetic reconstruction on the generation of mitochondrial genetic data from natural history collections. DNA extracted from toepad tissue allows mitochondrial genome assembly at relatively low sequencing depth compared to DNA from nucleated avian blood cells, as the latter contain significantly fewer mitochondria. While blood samples can be readily extracted from live birds, museum specimens offered a better opportunity for destructive tissue sampling. Therefore, a series of Galápagos Rail specimens deposited at the California Academy of Sciences (CAS) collected on Santa Cruz island in 1905– 1906 were accessed and toe pads sections from ten individuals were loaned. Additionally, modern samples (blood samples) were obtained from several islands to complement the genetic assessment of Galápagos Rails. The field work was carried out in four islands (May–July, 2017) and we concentrated our efforts to the highlands (> 500 m). We sampled the islands of Santa Cruz, Santiago, Pinta, and Isabela. On each locality we used playbacks to confirm the presence of rails as well as to define their territories. Birds were captured using the V-netting with playback trapping method [38], which consists in the arrangement of mist-nets forming a “V” and placed at ground level. Birds were lured inside the “V” using playback and led into the mist-nets by two people that monitor and adjust dynamically to bird responses. From each captured bird, we collected blood samples from the brachial vein in the field and blood was preserved on FTA blood cards (Whatman®).

### 2.2. Ancient DNA extraction and mitochondrial genomes

Ancient DNA from museum samples (toe pads) was performed at UCLA’s special ancient DNA facility following phenol–chloroform extraction procedures, but with 30 ul DTT added to the initial incubation step for the extraction from blood. DNA quantification was done using Epoch® (Bio Tek, USA) before library preparations and amplification procedures. DNA quality was tested using Qubit. Whole-genome next generation sequencing libraries with paired-end barcodes were prepared from five museum samples at the UCLA Technology Center for Genomics and Bioinformatics (TCGB). The samples were sequenced on the HiSeq3000 (150 PE) at the TCGB, yielding 2.13-2.37×10^7^ shotgun sequencing reads per sample. Read quality was determined using MultiQC [39]. There was little need for trimming, as reads were overall considerably shorter than 150 bp due to fragmentation.

Exploration and preliminary mapping to Swinhoe’s Rail *Coturnicops exquisitus* (Genbank accession no. NC_012143) was performed for all samples in Geneious v. 10.2.6 [40] using their native mapper with greatly relaxed settings, using ‘custom sensitivity’ only allowing mapping of full read pairs at minimum mapping quality 7, maximum gaps per read 30%, maximum mismatches per read 35%, and maximum ambiguity 10. We allowed up to 25 iterations, in which the first round reads are mapped directly onto the reference, and the subsequent iterations onto the consensus of the previously mapping reads, as a way of bridging regions of high divergence. Based on read quality, read length, and mapping success, we selected the sample GR9 (ORN 274 CAS catalog number) for a reference assembly of *L. spilonota*. For GR9, the mapping was complete in 5 iterations and 31,220 read pairs covered all of the *C. exquisitus* reference sequence. We extracted these as *fastq* files, which we then used as mitochondrion-enriched starting material for a *de novo* assembly and annotation of the mitochondrial genome with MitoZ v. 2.4.α [41]. Using three CPUs, we ran the *filter*, *assemble*, *findmitoscaf*, *annotate* and *visualize* modules with standard parameter settings, specifying --clade Chordata, --insert-size 200 and --filter_taxa_method 3. The resulting annotated genome was aligned with other rallid mitochondrial genomes and annotations were manually inspected and adjusted with regard to the start of 16S rRNA, and the includion of an additional C in ND3 causing a frameshift that is corrected through an unknown mechanism [42].

We mapped reads from the remaining samples to the circularized GR9 reference assembly in two steps with the Geneious mapper. First we enriched for mitochondrial reads in a step using the setting ‘low sensitivity’ with minimum mapping quality 15 and maximum mismatches per read 10% in two iterations. We then took the mapping reads and re-mapped them with the ‘highest sensitivity’ setting allowing up to 25 iterations. Resulting contig consensus sequences were aligned with the GR9 reference, all annotations lifted over, and every sequence variant position or ambiguity manually scrutinized against respective mapping reads.

### 2.3. Intraspecific genetics, genetic markers, and haplotype networks

In addition to the high-throughput sequencing described above, we sequenced two mitochondrial and one nuclear marker from a larger number of contemporary field samples. From each of the four visited islands we sampled 60 individuals in total to provide more detailed population structure and intraspecific patterns of diversification. Genomic DNA was extracted from blood cards using the Qiagen DNA Blood & Tissue Kit and following the manufacturer’s protocol. We focused on the mitochondrial genes cytochrome *b* (cyt*b*) gene and nicotinamide dehydrogenase 2 (ND2), and the nuclear recombination-activating gene 1 (RAG-1). Amplification and sequencing of cyt*b* was done using primers L14990–H16065 and primers L5143–H6313 for ND2 following the protocols described in Bonaccorso et al. [43], and for RAG-1, we used primers R52–R53 as described by Johansson et al. [44] (Table S1). Gel electrophoresis was used to confirm the presence and lenght of the amplifications, products were cleaned using ExoSAP-IT™ (Applied Biosystems™, Massachusetss, USA) and sequenced by Macrogen. Returning sequences were aligned, edited and trimmed using Geneious v. 10.2.6 [40]. Based on these sequences we represented intraspecific relationships from each locus using minimum-spanning networks using the package *pegas* (Populations and Evolutionary Genetics Analysis System) [45] as implemented in R 3.6.2 [46]. This package was also used to estimate haplotypic (*Hb*) and nucleotide diversity (p) as a whole group for each locus, as well as from individual islands with more than one haplotype.

### 2.4. Interspecific phylogenetic reconstruction of mitogenomes and divergence time estimates

#### 2.4.1. Coding mitogenome dataset

We followed Stervander et al. [47] and chose to analyse all coding regions (CDS) of the mitochondrial genome, partitioned per codon position (dataset “mtCDS”). As there was no appreciable phylogenetic signal between Galápagos Rail mitogenomes, we arbitrarily selected a single sample, GR5 (ORN 262 CAS catalog number), combined it through MAFFT alignent in Geneious to the dataset of Stervander et al. [47] and new extant gruiform mitogenomes (for taxa and accession numbers, see Figure 1a), resulting in a matrix comprising 32 species (22 species within Rallidae, 3 non-rallid species within Ralloidea, and 7 species within Gruoidea) and 11,418 base pairs (bp).

**Figure 1.**
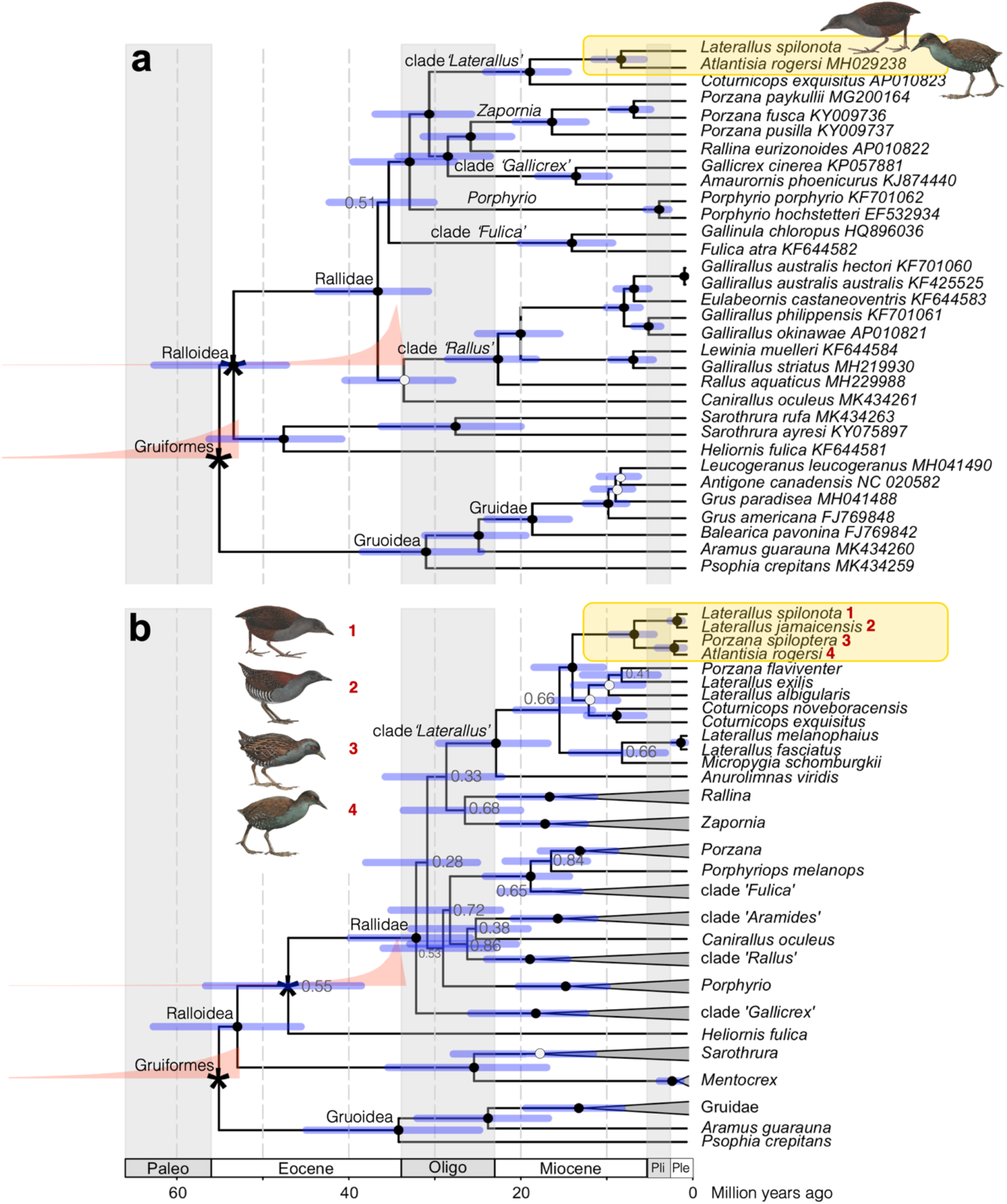
Dated phylogenies of family Rallidae showing the origin of the Galápagos Rail *Laterallus spilonota*, based on **(a)** all coding sequences of mitochondrial genes (dataset mtCDS; GenBank accession numbers stated at tips), and **(b)** the mitochondrial genes cytochrome *b* and cytochrome oxidase I and coding sequence of the nuclear recombination-activating gene 1 (dataset 2mt1nc; for GenBank accession numbers and taxon selection, see Data Accessibility). The chronograms are based on a relaxed clock model with fossil-based calibrations of stem Rallidae and crown Gruiformes. We also ran alternative analyses in which the Rallidae calibration was applied to the crown node, with yielding small differences of estimated node ages (see Table 2). Nodes that were time-calibrated are indicated with an asterisk, and the prior densities are drawn in salmon. Posterior probabilities (PP) are represented by filled circles (PP = 1.0), open circles (0.94 ≤ PP ≤ 0.99), or stated at nodes if lower. Node are drawn at median ages, with the 95% highest posterior density represented with blue bars. Shading represent geological periods—Paleogene (Paleo); Eocene; Oligocene (Oligo)—and epochs of the Neogene period: Miocene; Pliocene (Pli); Pleistocene (Ple). Every ten million years is indicated with dashed vertical lines. Illustrations reproduced with permission from Lynx Edicions ©.

We set up Bayesian Markov Chain Monte Carlo (MCMC) analyses for Beast v. 2.6.3 [48] partitioning per codon position across all mitochondrial genes [47], each fitted with a general time-reversible (GTR) model with four gamma categories (Γ) and an estimated proportion of invariant sites (I). We applied a relaxed log-normal clock model, let speciation follow a birth–death prior, constrained Ralloidea as monphyletic, and applied two priors for dating. Following Claramunt & Cracraft [49], we set a prior for crown Gruiformes (i.e. the root of the tree) following an exponential distribution with rate parameter 8.5 and offset 52 million years, that was fitted from the oldest fossils records from different parts of the world. We used the fossil data compiled in Stervander et al. [47], setting a prior for Rallidae following a lognormal distribution with mean 1.1, standard deviation 1.8, and offset 32.6 million years. However, as rightly pointed out by Garcia et al. [50], the inclusion of *Belgirallus* to calibrate crown Rallidae may not be correct, as *Belgirallus* may rather be representative of a stem rallid species [51]. We therefore primarily applied this prior to date stem Rallidae (i.e. parental node of Rallidae, being the Ralloidea node, the most recent common ancestor (MRCA) of Rallidae, Heliornithidae and Sarothruridae), and secondarily replicated the analyses applying this prior to crown Rallidae (i.e. the MRCA of extant Rallidae species). The rate scaler operators for A–C and C–T substitutions were modified (weight increased from 0.1 to 0.3) for improved performance.

We ran three replicates of each analysis for 75×10^6^ generations, sampled every 5×10^3^ generations, of which the first 5% were discarded as burn-in. Stationarity, high effectives sample sizes (ESS > 200), and between-replicate convergence were observed for almost all parameters (with the exception of some transition/transversion rate parameters for the second codon position, with ESS 135–200) in TRACER v1.4 [52], and maximum clade credibility trees with divergence times for both stem and root were obtained with median node heights calculated with TreeAnnotator [53]. Given convergence we used a single tree per analysis, and drew them in R 3.6.2 [46] using the packages ape v. 5.3 [54,55] and Phytools v. 0.6-99 [56].

#### 2.4.1. Two mitochondrial and one nuclear gene

In order to include more taxa, particularly focusing on the ‘American black rail clade’ *sensu* Stervander et al. [47], we created a dataset “2mt1nc” based on the two mitochondrial genes cyt*b* (1,068 bp) and cytochrome oxidase subunit I (COI; 747 bp), and the nuclear recombination activating gene 1 (RAG1; 930 bp). This dataset comprised 106 gruiform species, with varying degree of missing data. We followed the substitution model evaluation and partitioning by Stervander et al. [47], setting up Beast analyses partitioned by marker, with the substitution model HKY+Γ+I for and cyt*b* and COI, and K80+Γ+I for RAG1. Priors for clock model, speciation model, and calibrations followed those for the mtCDS dataset, as did operator modifications and run specifications, although number of generations 100×10^6^.

## 3. Results

### 3.1. Study sites and samples

We captured a total of 60 individuals in the field, 15 from Santa Cruz, 16 from Isabela, nine from Pinta, and 20 from Santiago. Out of the total number of samples from Santa Cruz, three were chicks, as were four from Santiago. Rails were found at sites above 500 meters characterized by high coverage of bracken (*Pteridium aquilinum*) and tall grass (*Pennisetum purpureum* and *Paspalum conjugatum*). Usually these sites were in a matrix with native Galápagos Miconia *Miconia robinsoniana* and invasive quinine trees *Cinchona pubescens* primarily in Santa Cruz, with guava trees *Psidium guajava* and blackberry (*Rubus* sp.) in the rest of the islands sampled.

### 3.2. Mitogenomes and genetic analysis

Sequencing produced an average 4.64×10^7^ (4.44–4.99×10^7^) reads per individual, with the average length per samples 58–76 bp due to fragmentation (Table S2). The complete GR9 reference *de novo* assembly produced by MitoZ was circularized and 17,045 bp long, with an average read depth of 61×. The GC content was 42.4% and it contained 13 protein-coding genes, two rRNAs, 22 tRNAs, and a 1,526 bp long control region (Figure S1). For the four remaining samples, the average number of reads per sample that mapped to the GR9 reference assembly was 81.9–345.7 (Table S2). Three samples (in addition to the reference GR9 also GR5 and GR8 (ORN 270 and 262 CAS catalog number respectively)) produced complete mitochondrial assemblies whereas two samples (GR2 and GR7; ORN 259 and 268 CAS catalog number repectively) contained ≤0.1% missing data.

The five mitogenomes contained 27 variants, comprising 23 single nucleotide polymorphisms (SNPs) and 4 insertion/deletion (indel) polymorphisms, distributed in protein-coding genes (15), control region (11), and 16S rRNA (1; Table S3). Out of the 15 SNPs in protein-coding genes, 4 were in codon position 1 and 11 in codon position 3, with 12 being synonymous mutation and 3 non-synonymous (Table S3). All variants were private to a single sample, with two exceptions: a synonymous SNP in COIII grouped GR7 and GR9 versus the three remaining samples, whereas an indel in the control region grouped GR9 and GR5 (reference haplotype), GR2 and GR8 (1 bp insertion), and GR7 (2 bp insertion; Table S3). Finally, there was mononucleotide length variation in the beginning of 16S rRNA which was ambiguous and unresolved due to low mapping success/coverage.

### 3.3. Interspecific phylogenetic reconstruction and divergence time estimates

The whole-mitochondrion phylogeny based on the mtCDS dataset, which did not include the Black Rail *Laterallus jamaicensis* or the Dot-winged Crake *L. [Porzana] spiloptera*, recovered the Galápagos Rail as a sister to the Inaccessible Island Rail *L. [Atlantisia] rogersi* with PP=1.0 (Figure 1a)

The mixed mitochondrial/nuclear marker phylogeny 2mt1nc, which contained more taxa but fewer base pairs, recovered the Galápagos Rail as sister to the Black Rail, the common ancestor of which was the sister of another sister species pair comprising the Inaccessible Island Rail and the Dot-winged Crake (Figure 1b). All nodes received full support with PP=1.0, and the estimated median age of the MRCA of the Galápagos Rail and the Black Rail was 1.1–1.2 Mya (95% HPDs 0.5–2.0 Mya), irrespective of whether the Rallidae prior was applied to the stem or crown (Table 1). The MRCA of all four species was estimated at a median age of 6.1 (stem) or 6.5 Mya (crown; for 95% HPDs, see Table 1), about a million years younger than the ages estimated from the mtCDS dataset.

**Table 1.**
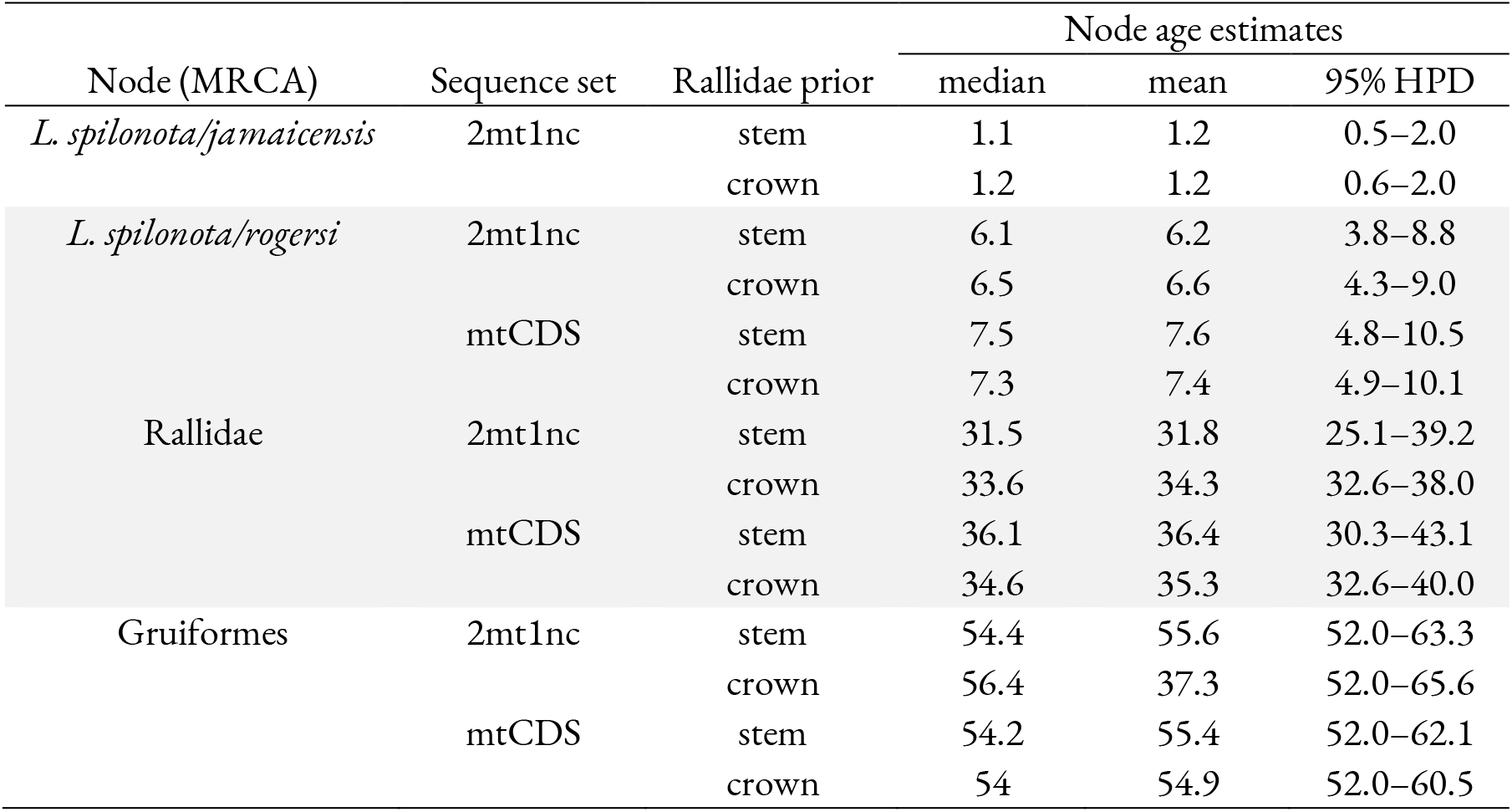
Estimated ages for relevant nodes (most recent common ancestors, MRCA) in the phylogenies created from the mtCDS and 2mt1nc sequence datasets, where the calibration density for the age of Rallidae was placed on either the crown or stem of the family. Age estimates are presented for the median and mean age as well as the 95% highest posterior density (HPD).

Overall, for both datasets, the placement of the Rallidae prior on the stem or crown had a small impact (≤9%) on the dating of both younger and older nodes, including the MRCA of Rallidae (median age 31.5–35.8 Mya; Table 1).

### 3.4. Intraspecific genetics, genetic markers, and haplotype networks

We obtained 59 sequences for cyt*b* (720 bp), 51 sequences for ND2 (969 bp), and 58 sequences for RAG-1 (302 bp). Overall genetic diversity (*Hb*) values were 0.439 for cyt*b*, 0.580 for ND2 and 0.034 for RAG-1. We report overall nucleotide diversity (p) of 0.0006 for cyt*b*, 0.0007 for ND2 and 0.0001 for RAG-1. Values for each island for both *Hb* and p are reported in Table 2, with Santa Cruz possessing the highest values for both *Hb* (0.791) and p for ND2 compared to the other islands and markers. Similarly, Santa Cruz and Santiago presented the highest values for both genetic metrics for cyt*b*. Pinta was the island that contained populations with the lowest values for all markers. RAG-1, being a nuclear marker, presented the lowest values across islands. Correspondingly, haplotype networks reported high levels of haplotype sharing across islands with shallow differences within haplotypes for each marker (Figure 2). We recovered three haplotypes for cyt*b*, all separated by 1 bp, shared across all islands; five haplotypes for ND2 comprising 4 bp difference, with no private haplotype at any given island; and two haplotypes for RAG-1 with 1 bp difference, one of the haplotypes private to Santiago. Cyt*b* haplotype I (LS02_Cytb) presented the highest frequency (n=43) and was found on all four islands, followed by haplotype II (LS07_Cytb; n=9) found on Santiago, Santa Cruz and Isabela, and haplotype III (LS06_Cytb; n=7) found on Santiago and Santa Cruz. ND2 haplotype I (LS05_ND2) had the highest frequency (n=32) and was found on all four islands, haplotype II (LS06_ND2; n=7) found only on Santiago and Santa Cruz, haplotype III (LS21_ND2; n=5) found only on Isabela, haplotype IV (LS03_ND2; n=4) found on Santiago and Santa Cruz, and finally haplotype V (LS07_ND2; n=3) found only on Santa Cruz. RAG-1 haplotype I (LS02_RAG1) presented the highest frequency (n=57) and was found on all four islands, whereas haplotype II (LS58_RAG1) was found in only one individual on Santiago (n=1).

**Figure 2.**
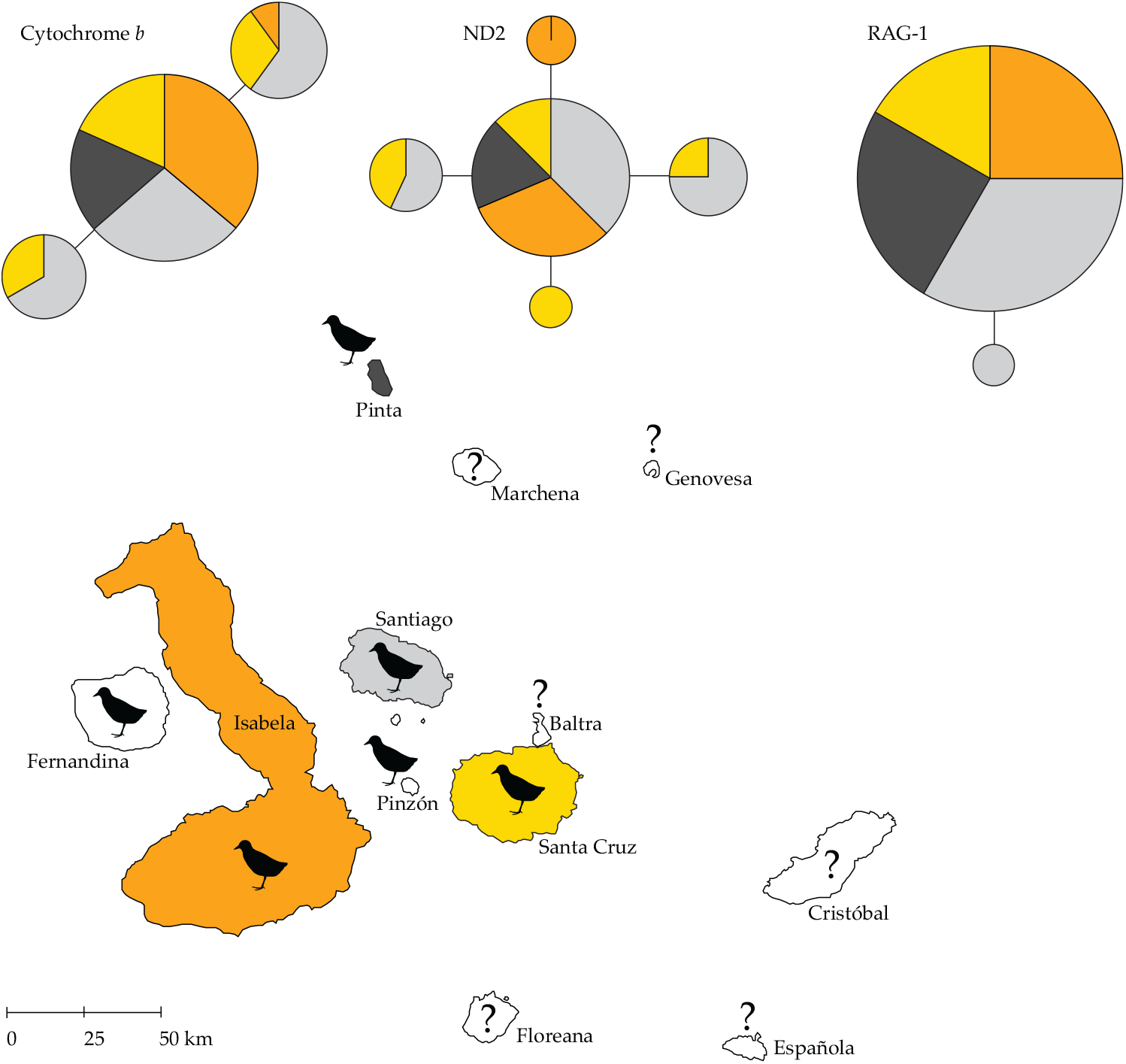
Haplotype network of mtDNA cyt*b*, ND2 and nuclear RAG-1 of *Laterallus spilonota* over a map of the Galápagos Islands. Each circle represents a different haplotype, the size of the circles is proportional to the number of individuals sharing that particular haplotype, and the color corresponds to the four islands analyzed here. Haplotypes differ from each other by one nucleotide represented by one branch. Islands with black rails represent the current distribution and question marks correspond to islands where rails are now extinct, or their presence is unknown.

## 4. Discussion

We found that the endemic Galápagos Rail forms a monophyletic group indicative of a single colonization event to the Galápagos Islands around 1.2 Mya, with negligible differences depending on whether the *Belgirallus* fossil was considered a crown or stem rallid for the calibration of the dated tree (Table 1), an overall pattern contrasting the findings of García et al. [50]. In comparison with other Galápagos land bird colonizers, rails coincide with the estimated arrival of Darwin’s finches (1.5–1.0 Mya [57,58] and flycatchers (genus *Pyrocephalus*: 1 Mya [59]; *Myiarchus*: 850 kya [60]). This also suggests that ancestors of the Galápagos Rail arrived to older islands first, most likely Cristóbal and Santa Cruz (~5 and 3 million years old respectively), moving west and colonizing new islands as these were formed by volcanic activity (Isabela ~0.5 Mya and Fernandina ~0.4 Mya, which it currently inhabits). Interestingly, speciation events have been idiosyncratic and variable in groups of the same time of origin. Finches have diversified into over 18 species [57] and *Pyrocephalus* flycatchers into two [59] compared to a single taxa of Galápagos Flycatcher *Myiarchus magnirostris* and Galápagos Rail. This phenomenon could be explained by an intrinsic evolvability in some groups compared to others, as explained by Chaves et al. [61] for Darwin’s finches and Hawaiian honeycreepers compared to other insular taxa of the same age, and sympatric to the same island systems.

Our phylogenetic reconstruction places the Galápagos Rail as sister to the Black Rail, confirming the suspicions by Leck [62]. Our findings also confirm the taxonomic placement for both *Laterallus* species, together with the Inaccessible Island Rail and Dot-winged Crake, within a clade defined as the “American black rails” with striking similarities in plumage coloration [31,47,63]. Additionally, and following island biogeographic theory [64], we suspected sister relationships of species to be closer if their distributions are also geographically close. In the case of the Galápagos avifauna, there are several examples of sister relationships between land bird species on the islands to continental species (Central America: Galápagos Yellow Warbler *Setophaga petechia aureola* [61], North America: Galápagos Hawk *Buteo galapagoensis* [65], North/Central America: Galápagos Flycatcher [60], South/North America: *Pyrocephalus* flycatchers [59]) and Caribbean forms (Darwin’s finches [66], Galápagos mockingbirds genus: *Mimus* [10]). The Black Rail presents a patchy distribution with resident subspecies populations on both the east (*L. j. jamaicensis*) and west (*L. j. coturniculus*) coast of North America (mainly US), Caribbean and Central America, with non-breeding (migrant) populations wintering in the Gulf of Mexico and Caribbean (*L. j. jamaicensis*) [67]. Another group of subspecies is distributed along the Pacific coast of Perú (*L. j. murivagans*) and Chile (*L. j. salinasi*), and from high elevation marshes and lakes around Lago Junín (*L. j. tuerosi*) at 4,200 m elevation in the central Andes of Perú [68,69]. It is likely that this species contains several phylogenetically distinct groups given the large, geographically-disjunct distribution and defined migratory behavior, and *L. j. tuerosi* has indeed been suggested to represent a distinct species [63,70]. Leck’s suggestion for the origin of the Galápagos Rail as derived from former migrants of North American Black Rail populations (*L. j. jamaicensis*) needs further exploration by including phylogenetic affinities between all Black Rail populations to pinpoint a possible origin of the Galápagos Rail. If proven right, it would not be the first group of migratory birds to have reached the Galápagos Islands, followed by the loss of this behavior to become a resident (and endemic) species of the archipelago (e.g. *Pyrocephalus* flycatchers, Galápagos Yellow Warbler).

The little genetic differentiation recovered from rail populations between islands is nevertheless surprising, particularly given the natural history of the species and time since colonization. Most insular rails are endemic to one island (or set of islands), suggesting a limited vagility after arrival [47,71,72]. Contrary to these notions, our haplotype reconstruction suggests high degrees of connectivity between islands. The absence of genetic structure in the three markers could be attributed to frequent movements among the islands that are on average 25 km apart, with the largest distance to Pinta 75–90 km from neighboring islands containing rail populations. These results were contrary to our prediction that Galápagos Rails should show higher levels of genetic structure not only based on its limited dispersal ability, but also given its habitat specialization. Rails are restricted to patchy marshes and meadows in the highlands and thus potentially support smaller local population sizes. It is important to mention that in the past, rails have been seen foraging near the coast in mangrove habitat [31], possibly increasing the chances to access to open water. It is possible then, that rails perform seasonal (or random) elevational migrations towards the coast and thus increase the chance for between-island connectivity as shown by the genetic sharing of haplotypes. A degree of swimming capacity reported for this species could explain this pattern [73], but alternative means of dispersal (i.e. rafting, nocturnal flights) are mere speculations at this moment.

Low levels of genetic diversity reported in Galápagos Rails could be indicative of the negative effects of past population bottlenecks. Galápagos Rails have suffered dramatically from the introduction of invasive species, either directly or indirectly. Goats were introduced after the first human settlements on the islands, with the largest impact in the 1960s and 1970s. Grazing goats stripped bare large expanses of native highland habitat, crucial for the presence of Galápagos Rails and other endemic fauna (giant tortoises). This ecological erosion combined with the presence of rats and cats, probably extirpated populations on Floreana, Baltra, Cristóbal, and Pinta islands [31,73,74] and most likely dramatically reduced populations numbers on surviving islands, as our homogenous genetic data indicates. After aggressive goat eradication programs and pest control programs were put in place, many of these islands became goat free (currently Isabela, Pinta, and Santiago). But not all islands suffered the impact of mammal introduction at the same magnitude. Varying levels of genetic diversity found across islands could be the result of such idiosyncratic historical events shaping rail demography on each island separately. For instance, Pinta populations showed the lowest diversity compared to other islands (mtDNA markers). Pinta was reported to be practically rail free after the introduction of goats [75], with a swift recovery only a few years after the start of eradication programs in late 1971 [73]. Currently rails are commonly found in high numbers thriving in the now recovered habitat. These past events could have left a genetic signature in present-day individuals, descendants from a reduced genetic pool of surviving birds. Alternatively, a complete extirpation on Pinta followed by a recolonization event from neighboring islands (founder effect) could also result in this pattern. The lack of private haplotypes on Pinta and the sharing of common haplotypes across islands supports the notion of high connectivity following island bottlenecks (or local extinction) and possible dynamic across-island recolonization events.

Natural history collections have served as a source of invaluable material to explore changes in declining, extinct or inaccessible taxa. Here, we relied on historical samples collected by the California Academy of Sciences in 1905–1906 to evaluate, for the first time, the phylogenetic reconstruction of the endemic Galápagos Rail. The combination of historical and modern samples allowed us to put forward a glimpse of the genetic history for this species and explore critical aspects of its evolutionary history. Unfortunately, several islands lost their rail populations in the last few decades (Floreana, Cristóbal, Baltra) and the magnitude of the historical loss of genetic variation could be great. Museum specimens collected prior to this ecological collapse (including from islands on which they are now extinct) could be used to inform the magnitude of the effect of invasive species on endemic Galápagos species. Genetic data produced from both present and historical samples has the potential to guide *in situ* management as well as translocations and reintroductions to islands currently with extirpated populations. The overall vulnerability of Galápagos Rail populations on the Galápagos is mirrored by the precarious conservation status of most insular rails around the world. Because the Galápagos Rail is inconspicuous, and one of the least studied land bird species on the Galápagos, its decline and extinction could go unnoticed. The implementation of our suggested framework in conjunction with the Galápagos National Park recovery/reintroduction program could directly benefit this endemic species and change the historical extinction record of flightless rails.

## Supplementary Information

The following are available as supplementary Information (pdf): Figure S1: Graphic representation of mitochondrial genome assembly, Table S1: Primer information, Table S2: Sequencing characteristics of mitochondrial genome, Table S3: Mitogenome sequence variation among museum samples, Table S4: Sequence information for mitochondrial cyt*b* and ND2, and nuclear RAG-1 markers.

## Author Contributions

Conceptualization, J.A.C.; methodology, J.A.C., E.A.D., S.E.U., A.C.B., and M.S.; software, J.A.C., P.J.M.T., S.E.U., A.C.B., and M.S.; validation, J.A.C., P.J.M.T., S.E.U., J.G.L., A.C.B., E.A.D. and M.S.; formal analysis, JA.C., P.J.M.T., S.E.U., A.C.B., and M.S.; investigation, J.A.C., P.J.M.T., S.E.U., J.G.L., A.C.B., E.A.D. and M.S.; resources, J.A.C., S.E.U., A.C.B., and M.S.; data curation, J.A.C., A.C.B., and M.S.; writing—original draft preparation, J.A.C.; writing—review and editing, J.A.C., P.J.M.T., S.E.U., J.G.L., A.C.B., E.A.D. and M.S.; visualization, J.A.C. and M.S.; supervision, J.A.C.; project administration, J.A.C.; funding acquisition, J.A.C. All authors have read and agreed to the published version of the manuscript.

## Acknowledgements

The authors want to thank the California Academy of Sciences (CAS) and its curator Maureen Flannery for granting us access to historical samples from Galápagos collected in 1905-06 expedition. To Matthew James for bringing to life the incredible story of the CAS expedition to the Galápagos and source of inspiration to this work. To Gabriela Gavilanez and Nathalia Valencia at the Laboratorio de Biología Evolutiva-USFQ. Special thanks to Dario F. Cueva for invaluable contribution towards data management. Permits to access genetic material in Galápagos were granted by Ministerio del Ambiente (MAE-DNB-CM-2016-0041) and Parque Nacional Galápagos (PC-63-18). Robert Wayne at UCLA provided critical support during ancient DNA extraction and sequencing.

## Data Accessibility

The new Galápagos Rail sequences (cyt*b*, ND2, RAG-1 alleles) have been deposited at GenBank with the accession numbers MW074873–MW074882 and mitochondrial reference genome with the accession number MW067132. Input and output files from the phylogenetic analyses have deposited at Zenodo: https://doi.org/10.5281/zenodo.4046620.

