## Supplemental Information for "Evolutionary history of the Galápagos Rail revealed by ancient mitogenomes and modern samples"

---

**Table S3**: Mitogenome sequence variation among five Galápagos Rail samples............................ pp 4–5
**Table S4**: Sequence information cyt*b*, ND2, and RAG-1 ........................................................... pp 6–7

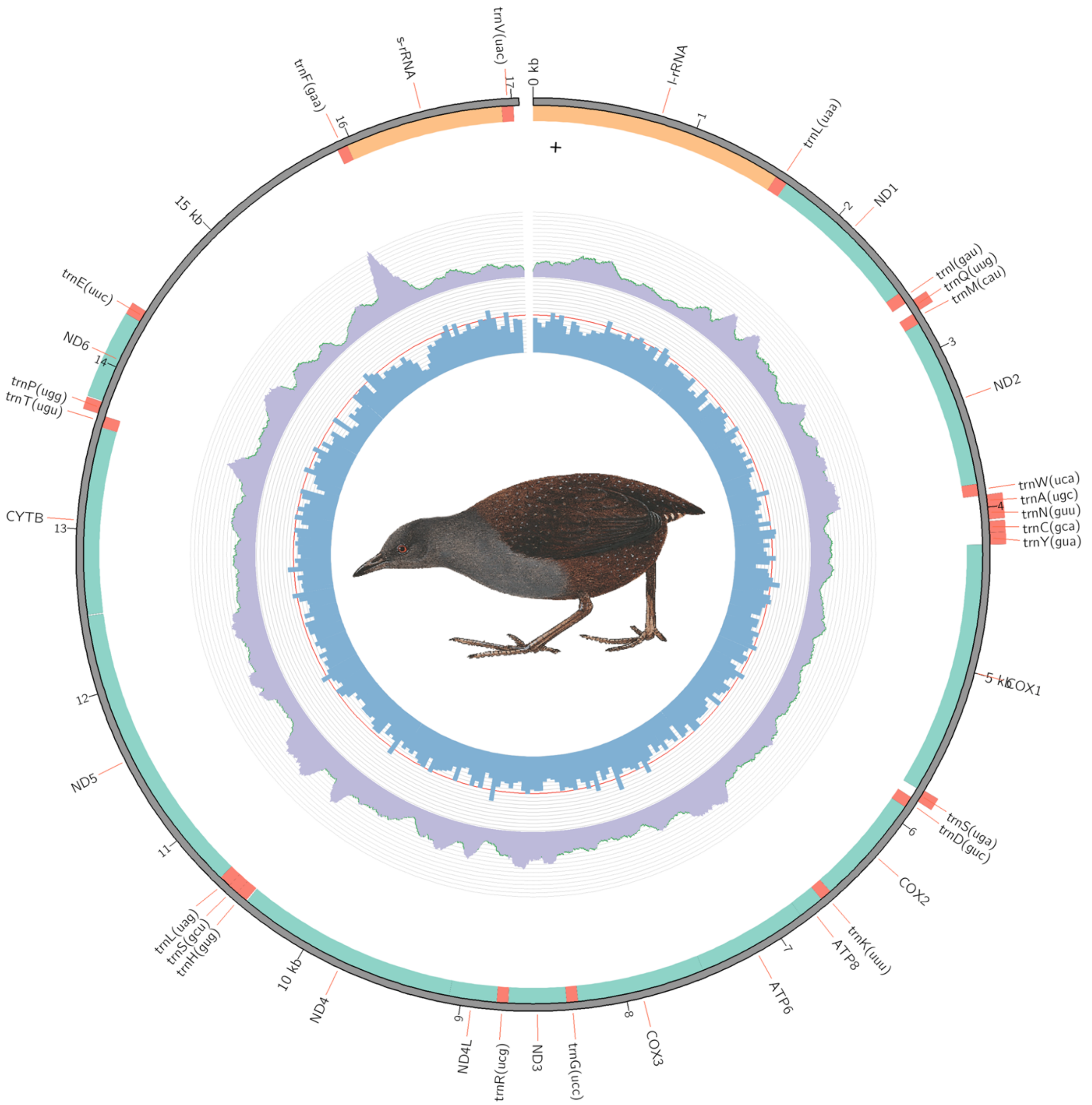

**Figure S1**. Graphic representation of the *Laterallus spilonota* reference mitochondrial genome assembly produced by MitoZ. The outer panel represents annotations of protein coding genes (teal), ribosomal RNAs (peach), and transfer RNAs (red); drawn on the inside for features on the +strand and on the outside for features on the –strand. The non-annotated region around 15 kbp is the control region (D-loop). Read coverage is drawn in grey in the middle panel, and GC content is represented by bluish bars in the in the inner panel. Illustration reproduced with permission from Lynx Edicions ©.

**Table S1.** Primer information

| Primer | Sequence (5′–3′) | Reference |
|---|---|---|
| **Cyt*b*** | | |
| L14990 | CCATCCAACATCTCAGCATGATGAAA | (Bonaccorso et al., 2010) |
| H16065 | GGAGTCTTCAGTCTCTGGTTTACAAGAC | (Bonaccorso et al., 2010) |
| **ND2** | | |
| L5143 | GAACCTACACARAAGRGATCAAAAC | (Bonaccorso et al., 2010) |
| H6313 | ACTCTTRTTTAAGGCTTTGAAGGC | (Bonaccorso et al., 2010) |
| **RAG-1** | | |
| R52 | CAAGCAGATGAAYTGGAGGC | (Johansson et. al., 2001) |
| R53 | TCCATGTCCTTTAAGGCACA | (Johansson et. al., 2001) |

**Table S2.** Sequencing characteristics of the five samples (first part of name), specified for read 1 (R1 in the last part of the sample name) and read 2 (R2). The Q20 and Q40 columns refer to proportion of bases of said quality or higher, with Q20 corresponding to a 99% and Q40 to a 99.99% base call accuracy. Number of reads mapped refers to the mapping onto the GR9 MitoZ reference *de novo* assembly.

| ***Sample name*** | **Number of reads** | **Duplicates** | **GC content** | **Mean length** | **Failed** | **≥Q20 (%)** | **≥Q40 (%)** | **N reads mapped** |
|---|---|---|---|---|---|---|---|---|
| *GR2_S8_R1* | 49,940,830 | 15.40% | 43% | 62 bp | 27% | 94.8 | 79.7 | 16,488 |
| *GR2_S8_R2* | | 14.90% | 43% | 62 bp | 36% | | | |
| *GR5_S9_R1* | 45,265,162 | 15.60% | 44% | 58 bp | 27% | 94.4 | 79.4 | 22,466 |
| *GR5_S9_R2* | | 14.80% | 44% | 59 bp | 36% | | | |
| *GR7_S10_R1* | 45,841,624 | 14.80% | 43% | 63 bp | 27% | 95.0 | 80.1 | 14,796 |
| *GR7_S10_R2* | | 14.10% | 43% | 64 bp | 36% | | | |
| *GR8_S11_R1* | 44,396,890 | 14.60% | 43% | 71 bp | 18% | 96.2 | 81.7 | 57,476 |
| *GR8_S11_R2* | | 14.20% | 43% | 72 bp | 18% | | | |
| *GR9_S12_R1* | 46,749,420 | 19.90% | 41% | 76 bp | 27% | 94.1 | 80.3 | 59,396 |
| *GR9_S12_R2* | | 19.60% | 41% | 76 bp | 27% | | | |

**Table S3.** Mitogenome sequence variation among five samples of the Galápagos Rail *Laterallus spilonota*. Position is indicated relative to the first base pair of the ND1 gene in the reference assembly of sample GR9 (GenBank accession no. MW067132). The last columns indicates in which sample a private allele occurs, length of insertions (+) or deletions (–), and/or the grouping of samples for phylogenetically informative variants.

| Variation type | Position | Unit | Codon position | Amino acid change | Informative | Sample / grouping |
|---|---|---|---|---|---|---|
| SNP | 285 | ND1 | 3 | synonymous | private | GR7 |
| SNP | 2,126 | ND2 | 1 | non-synonymous | private | GR9 |
| SNP | 3,716 | COI | 3 | synonymous | private | GR5 |
| SNP | 5,726 | ATP6 | 1 | non-synonymous | private | GR8 |
| SNP | 6,039 | COIII | 3 | synonymous | phylo-genetically informative | (GR7,GR9) \| (GR2,GR5,GR8) |
| SNP | 6,384 | COIII | 3 | synonymous | private | GR5 |
| SNP | 7,716 | ND4 | 3 | synonymous | private | GR7 |
| SNP | 7,833 | ND4 | 3 | synonymous | private | GR2 |
| SNP | 7,980 | ND4 | 3 | synonymous | private | GR7 |
| SNP | 8,622 | ND4 | 3 | synonymous | private | GR5 |
| SNP | 9,146 | ND5 | 3 | synonymous | private | GR7 |
| SNP | 9,344 | ND5 | 3 | synonymous | private | GR7 |
| SNP | 9,971 | ND5 | 3 | synonymous | private | GR8 |
| SNP | 11,808 | CYTB | 1 | non-synonymous | private | GR8 |
| Indel | 12,765 | control region | | | private | GR9 –1bp |
| SNP | 12,901 | control region | | | private | GR2 |
| SNP | 12,909 | control region | | | private | GR2 |
| SNP | 13,189 | control region | | | private | GR8 |
| SNP | 13,369 | control region | | | private | GR7 |
| SNP | 13,660 | control region | | | private | GR9 |
| SNP | 13,776 | control region | | | private | GR5 |
| Indel | 13,839 | control region | | | private | GR7 +1bp |
| SNP | 13,997 | control region | | | private | GR9 |
| SNP | 14,022 | control region | | | private | GR9 |

| | | | | | |
|---|---|---|---|---|---|
| Indel | 14,244 | control region | | phylo-genetically informative | (GR5+GR9) \| (GR2+GR8) +1bp \| GR7 +2bp |
| Indel | 15,401 | 16S rRNA | | ? | Ambiguous mononucleotide length variation |

**Table S4.** Sequence information for mitochondrial markers cyt*b* (A), ND2 (B) and nuclear RAG-1 (C). Haplotype name, frequency, polymorphic sites at given nucleotide position, sample ID, island, GenBank accession number and Sequence ID as submitted to GenBank.

**A) Cytochrome *b***

| Haplotype | Freq | Nucleotide position | | Individuals | Island | GenBank accession number | Sequence ID |
|---|---|---|---|---|---|---|---|
| | | 450 | 535 | | | | |
| LS02 | 43 | C | G | LS02 LS03 LS04 LS08 LS09 LS12 LS13 LS14 LS17 LS18 LS19 LS20 LS21 LS22 LS23 LS24 LS25 LS27 LS28 LS29 LS30 LS31_XX LS32 LS33 LS34 LS35 LS36 LS37 LS38 LS39 LS40 LS42 LS43 LS45 LS46 LS47 LS51 LS52 LS54 LS57 LS58 LS60 LSNN | Sta. Cruz Isabela Pinta Santiago | MW074873 | LS02_Cytb |
| LS06 | 7 | T | . | LS06 LS11 LS16 LS49 LS50 LS53 LS55 | Sta. Cruz Santiago | MW074874 | LS06_Cytb |
| LS07 | 9 | . | A | LS07 LS10 LS15 LS26 LS41 LS44 LS48 LS56 LS59 | Sta. Cruz Isabela Santiago | MW074875 | LS07_Cytb |

**B) ND2**

| Haplotype | Freq | Nucleotide position | | | | Individuals | Island | GenBank accession number | Sequence ID |
|---|---|---|---|---|---|---|---|---|---|
| | | 297 | 439 | 930 | 961 | | | | |
| LS05 | 32 | A | C | T | G | LS05 LS09 LS12 LS13 LS14 LS17S LS18S LS19S LS20S LS25S LS26S LS27S LS28S LS29S LS31_XX LS34 LS38 LS40 LS42 LS44 LS45 LS46 LS47 LS54 LS56 LS57 LS32- LS35 LS37 LS52 LS59 LSNNS | Sta. Cruz Isabela Pinta Santiago | MW074876 | LS05_ND2 |
| LS03 | 4 | . | . | . | A | LS03 LS04 LS08 LS60 | Sta. Cruz Santiago | MW074877 | LS03_ND2 |
| LS06 | 7 | . | T | . | . | LS06 LS11 LS16 LS50 LS53 LS55 LS49 | Sta. Cruz Santiago | MW074878 | LS06_ND2 |
| LS07 | 3 | . | . | G | . | LS07 LS10 LS15 | Sta. Cruz | MW074879 | LS07_ND2 |
| LS21 | 5 | G | . | . | . | LS21S LS22S LS23S LS24S LS30S | Isabela | MW074880 | LS21_ND2 |

**C) RAG-1**

| Haplotype | Freq | Nucleotide position | Individuals | Island | GenBank accession number | Sequence |
|---|---|---|---|---|---|---|
| | | 535 | | | | |
| LS02 | 57 | C | LS02 LS03 LS04 LS05 LS06 LS07 LS08 LS09 LS10 LS11 LS12 LS13 LS14 LS15 LS16 LS17 LS18 LS19 LS20 LS21 LS22 LS23 LS24 LS25 LS26 LS27 LS28 LS29 LS31 LS32 LS33 LS34 LS35 LSNN LS36 LS37 LS38 LS39 LS41 LS42 LS43 LS44 LS45 LS46 LS47 LS48 LS49 LS50 LS51 LS52 LS53 LS54 LS55 LS56 LS57 LS59 LS60 | Sta. Cruz Isabela Pinta Santiago | MW074881 | LS02_RAG1 |
| LS58 | 1 | G | LS58 | Santiago | MW074882 | LS58_RAG1 |
